# Subtle methodological variations substantially impact correlation test results in ecological time series

**DOI:** 10.1101/2024.10.11.617506

**Authors:** Caroline Cannistra, Linh Hoang, Alex E. Yuan, Wenying Shou

## Abstract

Correlation analyses using ecological time series can indicate phenomena such as interspecific interactions or an environmental factor that affects several populations. However, methodological choices in these analyses can significantly impact the results, potentially leading to spurious correlations or missed true associations. In this study, we explore how different decisions affect the performance of statistical tests for correlations between pairs of time series in simulated two-species ecosystems. We show that when performing nonparametric “surrogate data” tests, both the choice of statistic and the method of generating the null distribution can affect true positive and false positive rates. We also show how seemingly closely related methods of accounting for lagged correlation produce vastly different false positive rates. For methods that establish a null model by simulating the dynamics of one of the two species, we show that the choice of species simulated can influence test behavior. Additionally, we identify scenarios where the outcomes of analyses can be highly sensitive to the initial conditions of an ecosystem, even under simple mathematical models. Our results indicate the importance of thoughtful consideration and documentation of the statistical choices investigated here. To make this work broadly accessible, we include visual explanations of most methods tested in an appendix.

## Introduction

Research in ecology and environmental science frequently relies on the analysis of time series to discover interactions or shared influences among variables. However, this approach is subject to the well-known issue of spurious correlation. That is, a pair of time series might lack interactions or shared influences, but nevertheless appear to have a visually or numerically impressive correlation, such as the sizes of two populations that happen to both be growing during the data collection period [1, 2, 3, 4]. To avoid drawing misinformed conclusions from such spurious correlations, statistical tests designed to handle time series are essential.

A statistical test of correlation between a pair of time series typically involves three elements. First, one must select a null hypothesis, often that the two variables are independent (Box 1). Independence is a meaningful null hypothesis because disproving it indicates that one of the variables influences the other, or that they share a common causal influence [5, 6]. Second, one needs a correlation statistic (*ρ*) to measure a possible effect. Often a symmetric statistic is used in which *ρ*(*x, y*) = *ρ*(*y, x*) such as Pearson’s correlation coefficient [7, 8, 9], but some studies use an asymmetric statistic to suggest a directed relationship [10, 11, 12]. Third, determining the null distribution - how the correlation statistic behaves under the null hypothesis - is crucial (Fig 1). For certain correlation statistics, there may be an analytical expression for the null distribution (e.g. [7]). Alternatively, the null distribution may be estimated using surrogate data. To test for independence via this approach, one selects one of the variables to simulate – for instance *y* - and generates simulated copies of *y*, known as *y*\* surrogates, without using information from *x*. Then, the null distribution is estimated as the histogram of correlations between *x* and the various *y*\* surrogates [8, 13, 14].

**Figure 1:**
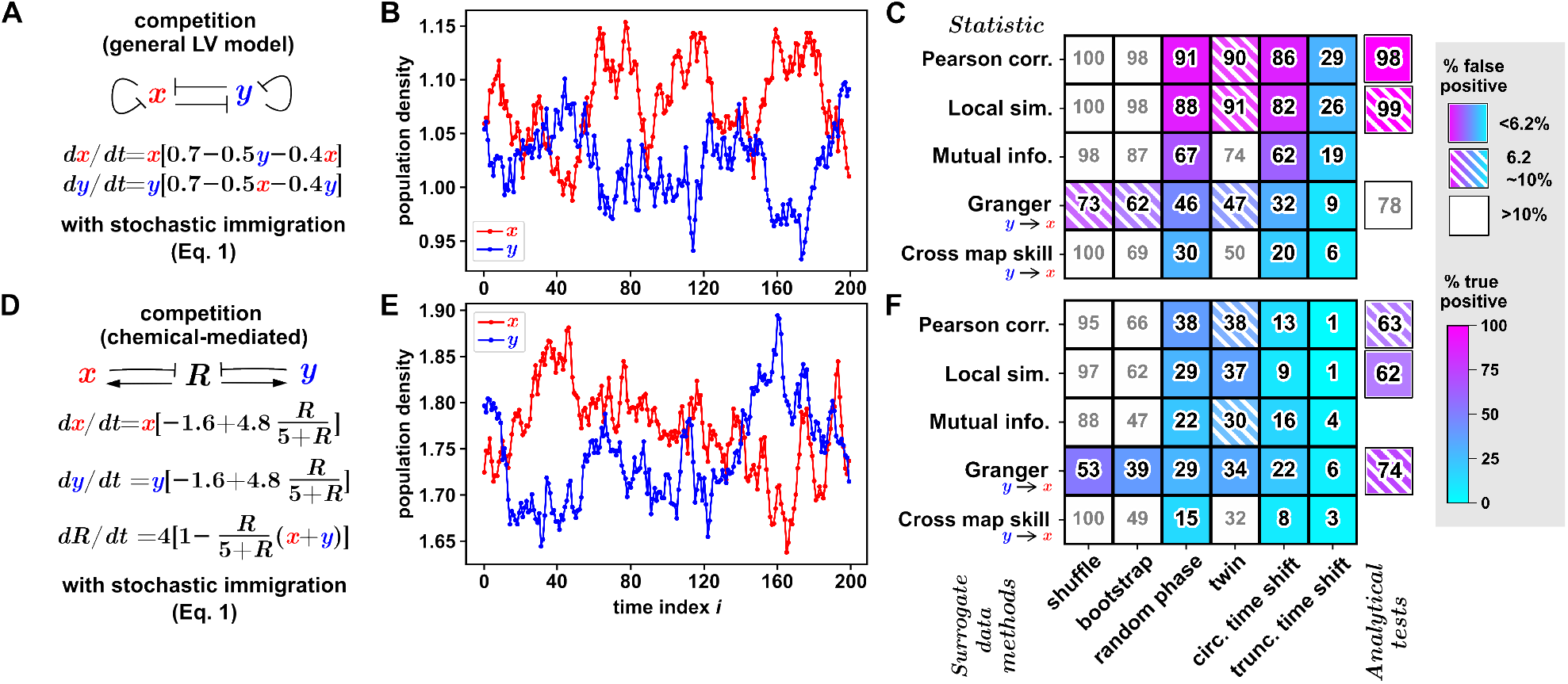
Choices of statistic and surrogate data method both affect test performance in two-species competition systems. (A, D): Models of competition. Two microbial species inhibit each other’s growth either directly (A) or indirectly through reduction of a shared resource *R* which is continuously added to the system (D). Populations are also subject to random fluctuations such as from stochastic immigration, modeled as noise terms (*ε*_*x,t*_ and *ε*_*y,t*_ in Eq. 1) drawn from a uniform distribution between 0 and 0.025 at regular intervals of step size Δ*t* = 0.05. The “measured” time series (B, E) to which dependence tests are applied have a lower sampling rate of once every five Δ*t* steps. (B, E): Negative correlation between population densities of competing species. (C, F): False positive and true positive rates when analyzing time series data using different combinations of statistic (rows) and surrogate data method or analytical test (columns). We show Granger causality and cross map skill in only one direction because the system is symmetrical and therefore results are the same in both directions. Solid shade, hatched shade, and white-out respectively represent ideal (*≤* 6.2%), questionable (between 6.2% and 10%), and unacceptable (*≥* 10) false positive rates. The number inside the box represents the percentage of true positives rounded to the nearest whole number, with a grey color used for tests with unacceptable false positive rates.

Given this grand buffet of methodological choices, it is no surprise that efforts to detect interspecies associations via time series analysis vary substantially between projects in their analysis pipelines. These differences, particularly when compounded with differences in experimental settings, can potentially contribute to inter-laboratory variation in results, even if multiple groups are studying the same system [15]. A better understanding of how these “devils in the details” impact results will empower researchers to better understand barriers to reproducibility and develop best practices for the field.

Here we explore this issue using a collection of two-species simulated ecosystems, with a focus on microbial communities. We report on four factors - other than the interaction network itself - that can impact the results of a correlation analysis. First, we show that both the correlation statistic and the method of generating the null distribution influence the true and false positive rates of tests. Second, we consider the issue of detecting correlations with a coupling lag, and we show that the true and false positive rates can vary widely depending on how a coupling lag is incorporated into the analysis. Third, for asymmetric correlation statistics, we show how initial conditions of an ecological system can influence the results of an association analysis in potentially surprising ways. Finally, in the context of surrogate data tests, we show that analysis results can vary substantially depending on which variable is chosen to be simulated.

### Box 1

#### Key Terminology

##### Dependence

Two events *A* and *B* are called independent if the probability that they co-occur is the same as the product of the probabilities that each occurs individually, i.e. *P*(*A* and *B*) = *P*(*A*)*P*(*B*). Likewise, two temporal processes *x* and *y* are called independent if the probability that *x* has some trajectory *x*′ and *y* has some trajectory *y*′ is the same as the product of the individual probabilities of these respective trajectories, i.e. *P*(*x* = *x*′ and *y* = *y*′) = *P*(*x* = *x*′)*P*(*y* = *y*′). Conversely, if there is some *x*′ and *y*′ such that this equation does not hold, then the processes are dependent. Dependence between two variables does not necessarily imply that one variable causes the other; two variables may be dependent if they share a common influence.

##### Independent and identically distributed

Random variables are “independent and identically distributed” (i.i.d.) if the possible state of each random variable is independent of the possible state of every other random variable, and if each random variable follows the same probability distribution. Data within a time series are usually not independent of each other because the past can influence the future.

##### Surrogate data

Artificial data generated from the observed data to simulate a null hypothesis. Surrogate data come with modeling assumptions, and violating the assumptions can lead to an invalid test (Table 2).

##### Stationarity

A stationary time series is one whose statistical properties do not change systematically over time. Formally, we call a time series (*x*_1_, …, *x*_*n*_) “stationary” if for any length *l < n*, the length-*l* subsequence (*x*_*i*_, *x*_*i*+1_, …, *x*_*i*+*l*−1_) is time-invariant, meaning that its statistical properties do not depend on *i*.

## Results

### Choice of both statistic and surrogate data method can affect test performance

We assembled a collection of tests for correlation and evaluated their true and false positive rates in simulated ecosystems consisting of two interacting species, whose population density values will be denoted by *x* and *y* respectively. To calculate true positive rates, we presented tests with data from the same simulation. To calculate false positive rates, we presented each test with the *x* time series from one simulation and the *y* time series from an identical but independent simulation of the same ecosystem. This is analogous to growing two replicate test tubes of the same microbial ecosystem and testing for a correlation between the *x* species of one tube and the *y* of another tube. In all cases, true and false positive rates were calculated as the percentage of detections (*p ≤* 0.05) reported out of 1000 total pairs of *x* and *y* time series.

We used 6 different methods for generating surrogate time series (Table 2) together with 5 different correlation statistics (Table 1), resulting in 6 × 5 = 30 combinatorially unique surrogate data testing procedures. See Appendix for a detailed introduction to each statistic and procedure. For three of the statistics we also included a parametric test (Fig 1C,F). All testing procedures are designed to test the null hypothesis that the two time series are independent, except for the parametric Granger causality test, which tests the null hypothesis of “Granger non-causality” (i.e. that the past of *x* does not contain unique information about the future of *y* [16, 17]). However, this exception does not matter for our purposes since all of our independent cases are also Granger non-causal, and all of our dependent cases are constructed to be Granger-causal in both directions.

**Table 1:** Statistics for measuring correlation between time series.

| Name | Description | Notes | Graphical explanation |
| --- | --- | --- | --- |
| Pearson's correlation | A commonly used statistic that measures global linear correlation. |  | N/A |
| Local similarity | A statistic based on sequence alignment algorithms that looks for temporally-local (i.e. transient) associations in time series [9]. |  | Appendix Fig. 1 |
| Mutual information (MI) | A statistic that measures nonlinear association [28, 29]. | When data are independent and identically distributed, MI has the nice property that $MI > 0$ if and only if two variables are dependent. However, MI estimators applied to autocorrelated data do not have this property [30]. | Appendix Fig. 4 |
| Granger causality (GC) | Consider a system of two variables $x$ and $y$. To compute Granger causality in the $x \rightarrow y$ direction, predict later values of $y$ using a linear model that includes earlier values of either (A) both $x$ and $y$, or (B) includes history of $y$ only; then Granger causality measures the difference in accuracy of the two models [27, 16, 31]. | Although GC is commonly used with a parametric test, tests based on the GC principle have been performed using surrogate data [32, 11]. | Appendix Fig. 2 |
| Cross map skill | Cross map skill in the $x \rightarrow y$ direction measures the amount of information about a potential causer variable ($x$) stored in the history of a potential causee variable ($y$)[10, 33]. | Cross map skill is the statistic used in “convergent cross mapping”, a technique that attempts to detect directed causal relationships in deterministic systems[10]. | Appendix Fig. 3 |

We first assessed how well these tests perform when given time series of population densities from a pair of species engaged in competition. We used stochastic simulations based on two underlying models: a Lotka-Volterra model [18, 19, 20] where each species’ growth equation includes an interaction term that directly encodes the negative impact of its competitor (Fig 1A), and a resource-mediated interaction model [21] where species compete via their shared need of a finite resource (1D).

False positive rates of surrogate data tests are sensitive to both the null model and the correlation statistic (Fig. 1C, F). Notably, the random shuffle and block bootstrap null models produce unacceptable false positive rates with most correlation statistics (except Granger causality). The poor performance of these null models is unsurprising since both scramble data in time, destroying the original autocorrelation to varying degrees. True positive rates are also sensitive to both the surrogate procedure and the correlation statistic.

We asked whether the ranking of correlation statistics by statistical power depends on the null model, and vice versa. In both the Lotka-Volterra and chemical interaction settings, the ranking of null models is identical across statistics with only one exception that was not statistically significant (*p >* 0.1, two-tailed z-test of proportions). In contrast the ranking of statistics can depend on the null model. For example, in the chemical interaction setting, mutual information has higher power than local similarity with circular and truncated time shift, but the reverse is true when twin or random phase surrogates are used. We recovered the same result (with *p <* 0.005) when repeating each of these 4 comparisons with a separate random number seed (to avoid the need to statistically account for 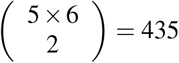 possible pairwise comparisons). Thus, to maximize power, a researcher may need to select their correlation statistic based on the null model they are using.

### Different methods of accounting for lagged correlation affect test performance

Often the effects of interactions only become apparent after a delay, leading to a pair of time series whose correlation is detectable only when one time series is shifted relative to the other by an appropriate lag. In this case, a method is required to choose a lag for both the original and surrogate time series. One way is to choose the lag that gives the highest correlation between the two original time series, and then use the same lag when constructing the null distribution with surrogate time series. We call this the “fixed lag” approach. We suspected that under the fixed lag approach, tests are likely to produce high false positive rates because the original time series enjoys an optimized lag, but the surrogates do not, resulting in an unfair null model (Fig. 2Bii-iii, condition *b*). Alternatively, one can optimize the lag for the original correlation and then independently optimize the lag for surrogate correlations (“tailored lags”). Since the tailored lag approach applies the same pre-processing steps to both the original and surrogate data, it seems more likely to produce valid tests (Fig. 2Bii-iii, condition *a*).

**Figure 2:**
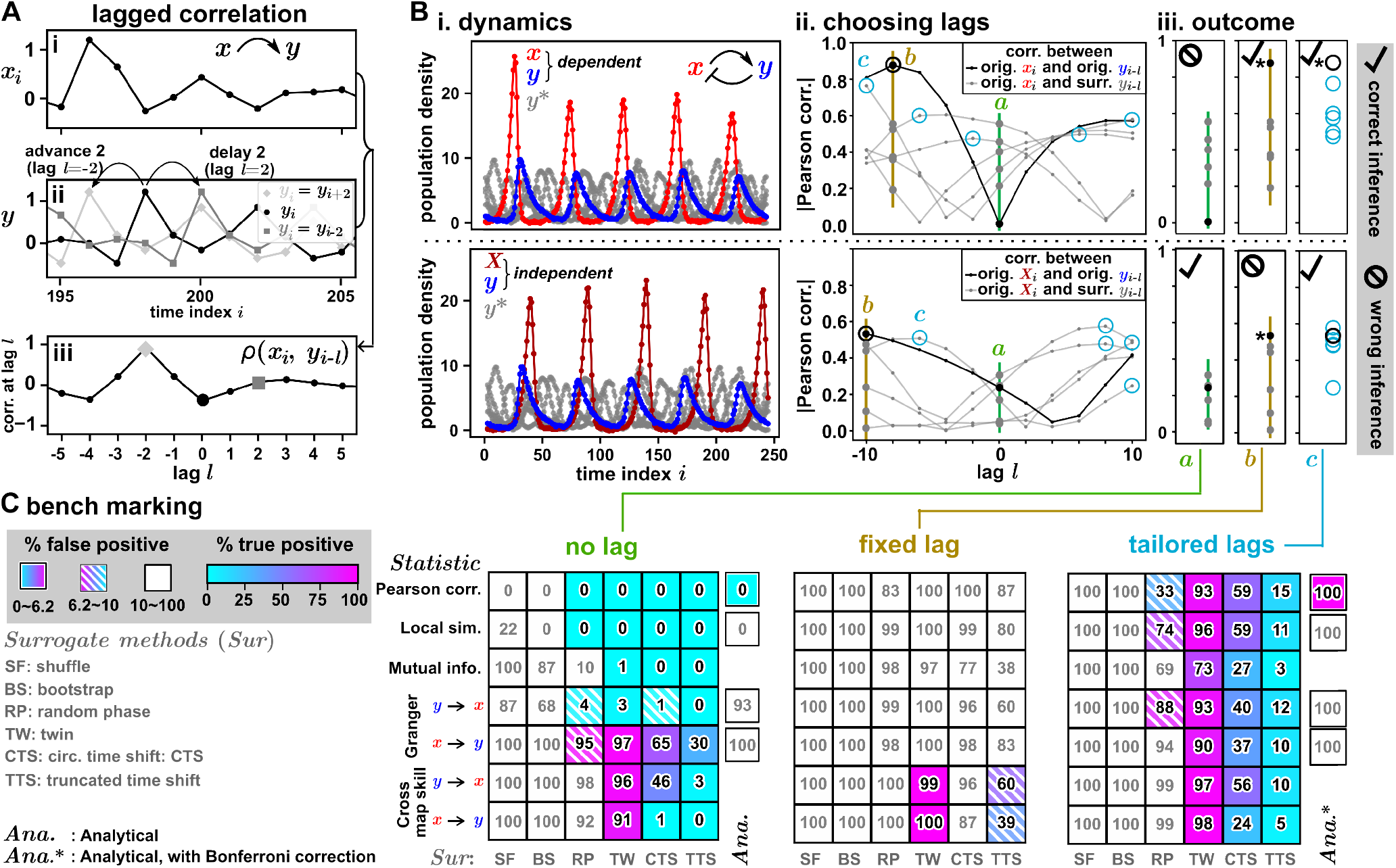
Different strategies of choosing a lag in the correlation function lead to substantially different test performance. (**A**) Lagged correlation may be required to reveal dependence with a delayed response. Here the *x* variable (Ai) influences the *y* variable (Aii) with a lag of 2. Aii depicts the original *y* series (black circles), the series shifted by *l* = −2 (light grey diamonds), and shifted by *l* = 2 (dark grey squares). The Pearson correlation between *x*_*i*_ and *y*_*i*−*l*_ is calculated at different time lags *l* (Aiii), peaking at lag *l* = −2 (light grey diamond). (**B**) Three strategies for choosing correlation lags. Bi illustrates time series from a Lotka-Volterra predator-prey system in which predator dynamics lag behind prey dynamics, with the top panel showing a predator (*y*; blue) and prey (*x*; red) from the same simulation and the bottom panel showing a predator (*y*) and prey (*X*; brown) from different simulations. Both panels share identical *y* series and random phase surrogates (*y*\*; grey). Bii depicts correlations between *x* and the original *y* calculated at different lags *l* (black curve), or between *x* and five representative surrogates *y*\* (grey curves). Shown are three ways to choose original and surrogate lagged correlations. With no lag (*a*, bold dots on green line), all correlations are calculated at lag *l* = 0. With a “fixed lag” (*b*, bold dots on gold line), all chosen correlations are calculated at the lag that maximises that of the original series. With “tailored lags” (*c*, black and cyan open circles), for each pair of time series, either original (black) or surrogates (cyan), the lag with the highest correlation is used. Biii shows the original and surrogate correlations under each lag method isolated on a line to clarify their ranking. An asterisk (*) indicates detection of a significant correlation with *p* = 1/6 (in practice, at least 19 surrogates are needed to achieve *p* = 0.05). A check mark indicates a correct inference (rejecting the null hypothesis when *x* and *y* are truly dependent, or failing to reject when *x* and *y* are truly independent);, incorrect inference is represented by a slashed circle. In this simplified example, the “no lag” approach always fails to detect a correlation, the “fixed lag” approach always detects a correlation, and the “tailored lag” approach produces correct results. (**C**) A simulation benchmark of the three strategies. Lags, when used, were chosen from between −10 and 10 in steps of size 2. Analytical tests were included in the “no lag” condition. We also tried applying analytical tests at each lag, reporting the lowest *p*-value and then performing a Bonferroni correction; this procedure’s results are shown next to the “tailored lags” table (*Ana*.* column). Interpretation of figure (C) is similar to 1. See Methods for simulation details.

We verify these expectations by applying the different lag strategies to detect dependence between the density of a predator and its prey in a predator-prey Lotka-Volterra simulation. When the two species are in the same system, predator dynamics lag behind prey dynamics. When no lags are used in the analysis (Fig 2C), tests mostly either produce unacceptable false positive rates, or very low true positive rates. The exceptions are tests using Granger causality and cross map skill–the two correlation methods among those we tested that necessarily incorporate lags into the calculation of the statistic. The fixed lag approach produced mostly unacceptable false positive rates, as expected. Finally, the tailored lag approach tended to result in broadly superior performance, with false positive rates similar to the no-lag condition, but with true positive rates generally exceeding those of the no-lag condition.

### Tests based on Granger causality and cross map skill are sensitive to system state

Microbial ecosystems often feature multiple steady states. That is, due to chance events, similar microbial communities can settle into distinct states characterized by unique species compositions. Here we show that, perhaps counterintuitively, the results of Granger causality or cross map skill analyses can be highly sensitive to the system’s state.

To illustrate, consider a two-species competitive microbial community that is symmetric in the sense that the equations describing their dynamics follow precisely the same form, with identical parameters for growth, self-inhibition, and cross-inhibition. This system has two stable states characterized by the dominance of one or the other species. We choose an initial condition in which the *x* species is more abundant than *y*. Consequently, *y* remains at high density with some minor fluctuations, while the *x* density remains near zero (Fig. 3A, top half). Since Granger causality and cross map skill are both directional statistics, we can think of applying these statistics in the *x → y* and *y → x* directions as applying them to the same system in different states.

**Figure 3:**
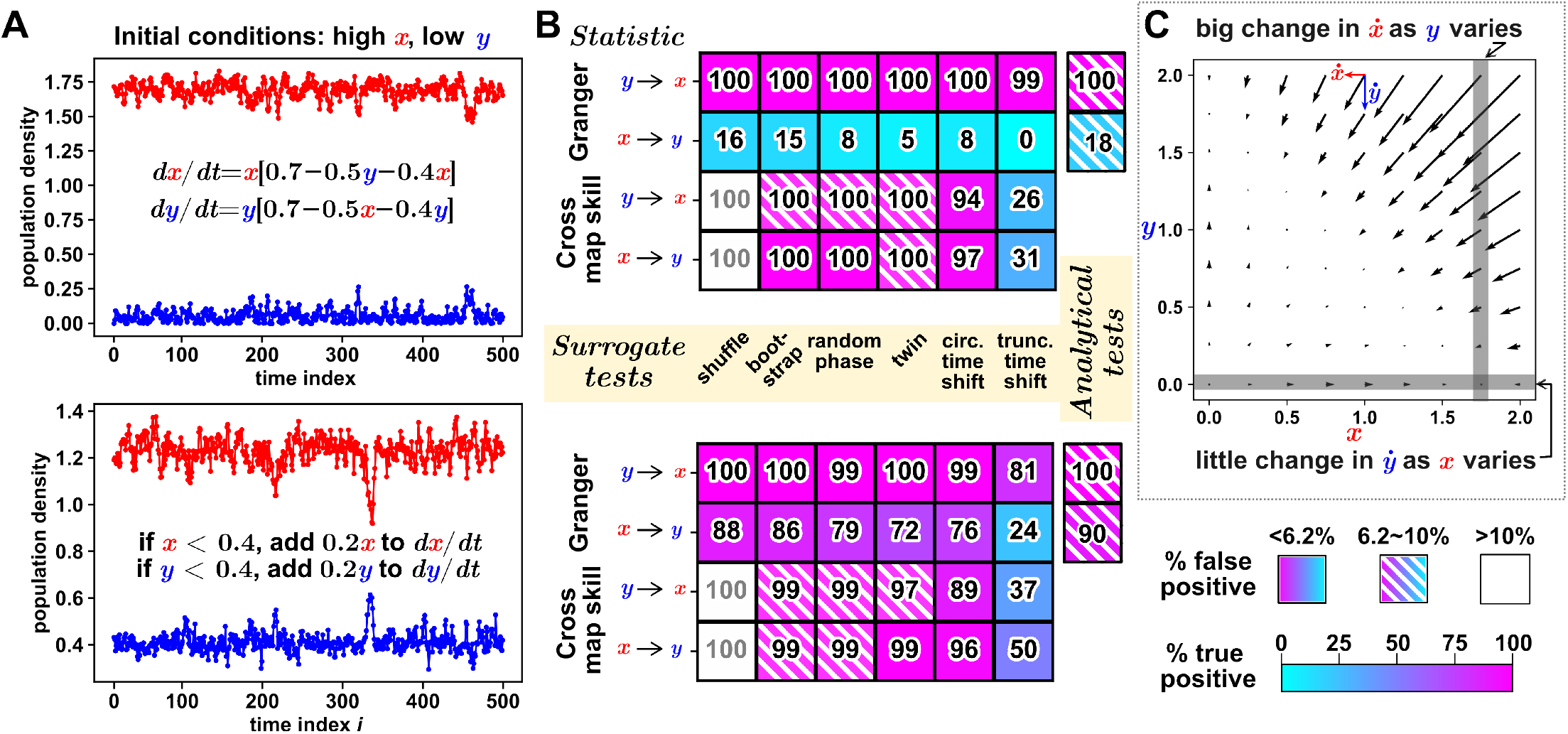
In multi-stable microbial communities, tests based on Granger causality and cross map skill can depend on community state. (**A**, top): A two-species competitive microbial community in which the two species (*x* and *y*) have identical parameters for growth, self-inhibition, and cross-inhibition. There are two steady states (dominance of one or the other species). The system is initialized in the state where *x* dominates. Both populations experience random fluctuations, so the suppressed species is not permanently extinct. (A, bottom): Similar to the top panel of (A), except here, if a species’ population density drops below 0.4, its intrinsic growth rate is increased by 0.2. (**B**): Surrogate data tests of dependence using Granger causality or cross map skill applied in either the *x → y* direction (corresponding roughly to the alternative hypothesis that *x* drives the dynamics of *y*), or in the *y → x* direction. Analytical Granger causality tests are also shown. The top and bottom tables correspond to the top and bottom panels shown in (A), respectively. (**C**): A vector field plot corresponding to the differential equations shown in the top half of (A), indicating that at the stable community state, changes in *y* lead to large changes in *dx*/*dt* (written as 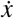 to conserve space), whereas the reverse does not hold. See Methods for simulation details.

Since the interaction parameters are equal, we might expect that tests using directional statistics would detect dependence regardless of the statistic’s direction. However, the tests that use Granger causality in the *y → x* direction enjoy statistical power close to 100%, whereas the same tests in the reverse direction have power below 20% (Fig. 3B, top half). This effect occurs in both the analytical Granger causality test (a genuine test of prediction improvement, i.e. “Granger causality”) and the surrogate data-based tests (which are technically tests of independence). One possible explanation for this asymmetry becomes apparent upon inspecting the vector field plot for the deterministic version of the system. Specifically, changes in the density of the suppressed species (*y*) have a relatively large effect on the dynamics of the dominant species (*x*), whereas the reverse does not hold (Fig 3C). Cross map skill is less sensitive to changes in the direction (Fig. 3B, top half;*x → y* versus *y → x*).

To determine whether this effect is entirely due to one species being at near-zero abundance, we also examine a system of two competing microbial species that coexist at different densities. Here, both species have a faster birth rate when their density falls below a threshold, so the lower-density species tends to stay near this threshold and away from zero (Fig. 3A-B, bottom half). Once again, the Granger causality test in the *y → x* direction achieves greater power, although the difference is less dramatic in this case. Moreover, in this setting, cross map skill can also exhibit modest sensitivity to the direction, achieving greater power in the *x → y* direction than *y → x* with circular time shift surrogates (96% versus 89%; *p <* 10^−8^) and truncated time shift surrogates (50% versus 37%; *p <* 10^−8^).

### Choice of surrogate template affects test performance

Surrogate data testing is most commonly implemented by replacing just one of the time series with surrogates, not both. This has the theoretical advantage that only one time series must satisfy the assumptions of the surrogates in order for the test to be valid. Yet, the decision of which time series to use as the surrogate template often appears arbitrary and a rationale is rarely given in practice.

Here we show that test performance can depend on which variable is used as the surrogate data template. We use as our ground-truth a two-species competition model similar to that of Fig 3A (but with a shorter sampling period; see Table 3) where *x* tends to dominate *y* due to a greater initial density. We assessed the performance of tests when either *x* or *y* was used as the surrogate data template (Fig 4C, “surrX” and “surrY” respectively).

**Table 2:** Surrogate data generation methods. Graphical explanations can be found in the appendix. ^*a*^A test is exactly valid under its modeling assumption if the following criterion is strictly satisfied: The false positive rate - the chance of erroneously reporting dependence - will be no more than the significance level *α* if we infer dependence only when the *p* value is equal to or less than *α*.

| Name | Summary of surrogate data generation procedure | Assumption about the time series used to produce surrogates | Is the test approximate or exactly valid<sup>a</sup> under its assumption? | Notes | Graphical explanation |
| --- | --- | --- | --- | --- | --- |
| Random shuffle / permutation | Shuffle all observations in time. This preserves distribution of the data points while destroying autocorrelation (correlations between time points within a series) [14]. | Time points are independent and identically distributed (i.i.d.) random variables. | Exactly valid [34, 35] | The i.i.d. assumption is rarely reasonable in time series. | Appendix Fig. 5A |
| Stationary bootstrap | Shuffle randomly sized “blocks” of consecutive data points in time [36, 32, 11]. Shuffling blocks instead of individual points preserves some of the original time series’ autocorrelation. | The time series is stationary. | Approximate: Junctions between blocks can create disruptions [37]. |  | Appendix Fig. 5B |
| Iterative amplitude-adjusted Fourier Transform (“random phase”) | Iterate between two steps. One step uses a Fourier Transform to ensure that a surrogate has the same power spectrum as the original data; the other step ensures that a surrogate has the same distribution as the original data. Over time, surrogates can be generated with accurate power spectra and distributions [38]. | The time series is obtained by taking a stationary multivariate Gaussian, and passing it through a possibly nonlinear time-independent function [38, 39]. | Approximate: Accuracy is found to be good but not perfect [38]. | Also used to test for so-called “nonlinear processes” [38, 39]. | Appendix Fig. 6 |
| Twin | Shuffle randomly sized “blocks” of consecutive data points such that the neighboring ends of adjacent blocks follow similar trajectories, limiting discontinuities between blocks [40]. | Unclear | N/A |  | Appendix Fig. 9 |
| Circular time shift | Shift all time points by the same amount, with the end of the time series looping back to the beginning. This preserves all temporal structure except at the ends, and may introduce a discontinuity between the ends [14, 13]. | The time series is stationary. | Approximate: Early and late times are artificially joined [37]. | Surrogates are not independent. | Appendix Fig. 8A |
| (Adjusted) Truncated time shift | $x$ is truncated so that it runs from $i_1$ to $i_2$. Each surrogate $y^*$ is then a subsequence of $y$ from $i_1 + \delta$ to $i_2 + \delta$, where $\delta$ is varied to produce different surrogates. The case of $\delta = 0$ is considered the “original” $y$ [13, 37]. | The time series used to generate surrogates is stationary. | Exactly valid with appropriate $p$-value adjustment [37] | Surrogates are not independent, but the $p$-value is adjusted to account for this, thus resulting in an exactly valid test. | Appendix Fig. 8B |

**Table 3:** Simulation parameters for each figure. Unif(*a, b*) and 𝒩(*μ, σ*^2^) denote a uniform distribution between *a* and *b* and a normal distribution with mean of *μ* and standard deviation of *σ* respectively. The integration step size Δ*t* is 0.05 for all systems.

| Figure | Number of time points | Sampling period | Process noise distribution (per integration step) | Measurement noise variance (per time point) | Initial conditions |
| --- | --- | --- | --- | --- | --- |
| 1A-C (direct competition) | 200 | $5\Delta t$ | $\text{Unif}(0, 0.5\Delta t)$ | 0 | $x, y \sim \text{Unif}(0, 5)$ |
| 1D-F (indirect competition) | 200 | $5\Delta t$ | $\text{Unif}(0, 0.5\Delta t)$ | 0 | $x, y \sim \text{Unif}(0, 5)$<br>$R = 0$ |
| 2 (predator-prey) | 250 | $5\Delta t$ | $\mathcal{N}(0, (0.2\Delta t)^2)$ | $10^{-2}$ | $x, y \sim \text{Unif}(0, 2)$ |
| 3A-C (competition with one extinct) | 500 | $25\Delta t$ | $\mathcal{N}(0, (0.2\Delta t)^2)$ | $10^{-6}$ | $x = 2, y = 0$ |
| 3D-F (competition with coexistence at different densities) | 500 | $25\Delta t$ | $\mathcal{N}(0, (0.2\Delta t)^2)$ | $10^{-6}$ | $x = 2, y = 0$ |
| 4 A-C (competition with one extinct) | 500 | $5\Delta t$ | $\mathcal{N}(0, (0.2\Delta t)^2)$ | $10^{-6}$ | $x = 2, y = 0$ |

**Figure 4:**
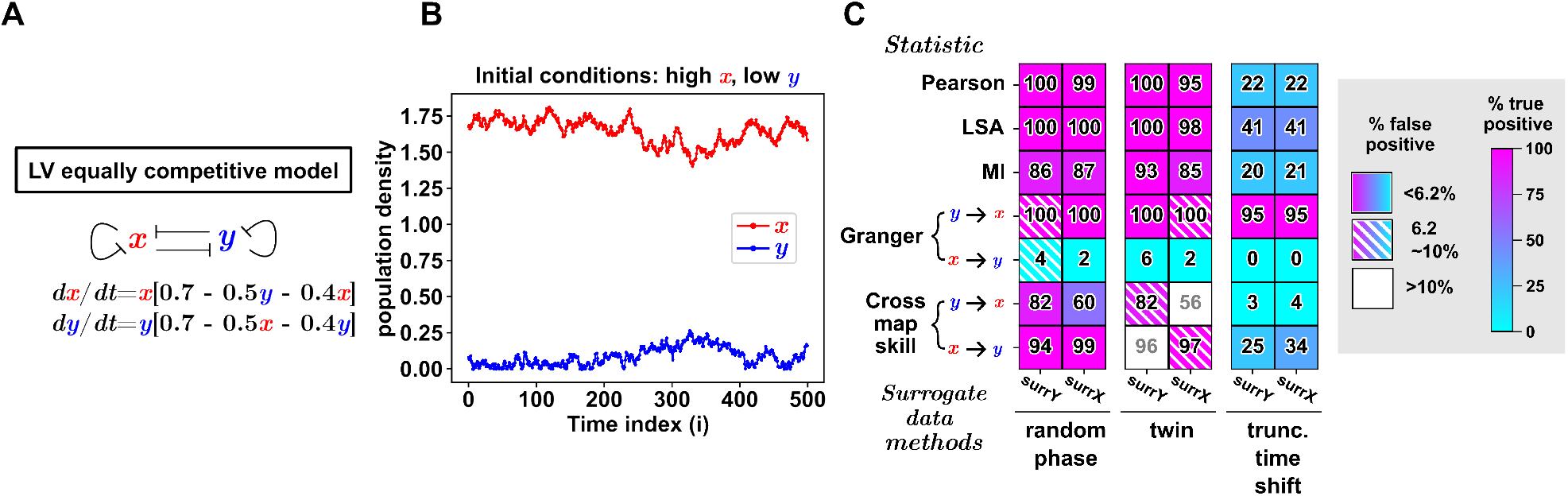
The behavior of surrogate tests differs between using surrogates *x*\* and surrogates *y*\*. (A) System equations used for simulation. (B) Example time series. Simulations begin with *x* at a greater population size than *y*, introducing an asymmetry wherein *x* is the dominant competitor. (C) Test performance. Each test is performed using either surrogates of *y* (surrY) or surrogates of *x* (surrX). Lags such as in Fig 2 were not used in these correlation tests. See Methods for simulation details.

The effect of surrogate template choice was particularly striking when twin surrogates were used with cross-map skill, a directed statistic commonly associated with a method that attempts to infer the direction of causality (Table 1). Producing surrogates from the presumed causee variable (Fig 4C, *y → x* with surrX or *x → y* with surrY) led to false positive rates above 20%. Conversely, when the presumed causer variable was used as the template (e.g. *y → x* with surrY), tests based on cross map skill and twin surrogates produced more modest false positive rates below 8%.

It is also possible for non-directional statistics to exhibit sensitivity to the choice of surrogate template. The twin surrogate test with mutual information as the statistic enjoyed greater statistical power when surrogate were generated from *y* than *x* (93% versus 85%, *p <* 10^−7^).

## Discussion

The numerical experiments presented here indicate that within a surrogate data test for correlation, there exist numerous consequential choices. Test performance is sensitive to both the correlation statistic and the null model, and the two elements interact in such a way that the preference of one statistic over another may depend on which null model is used (Fig 1F). Test results can depend on whether and how lags are included in the correlation analysis, with a naive method (Fig 2, “fixed lag”) producing unacceptably high false positive rates. Changing which variable is used as the surrogate template can impact test performance (Fig 4C). This effect was particularly striking in the case of a test using cross map skill as the statistic and twin surrogates as the null model, where using the presumed causing variable as the surrogate template produced a false positive rate under 8%, while producing surrogates from the presumed causee led to a false positive rate above 20%. Note that both the presumed causer and causee have been used as the surrogate template when testing the significance of cross map skill [22, 23, 24, 25].

Choices such as these represent opportunities for research results to reflect a scientist’s preferred statistical recipe, perhaps as much as they reflect the data. What is to be done? We offer two suggestions. First, when a choice presents itself (e.g. among correlation statistics, null models, or the choice of surrogate template), we recommend that researchers consider performing their analysis on at least two variations to assess whether and how the analysis conclusions are affected. Second, we advocate for greater transparency in testing methodology. For instance, introducing lags into correlation tests is commonplace, but it is not always clear whether studies are using the correct “tailored lags” approach or the naive “fixed lag” method. Similarly, It is routine to use surrogate data procedures to test the significance of a correlation, but it is often unclear which time series is used as the surrogate template.

Beyond the effects of decisions within statistical procedures, we illustrated that for microbial communities with multiple stable states, the community state can profoundly impact the results of an analysis (Fig 4). Thus, the correlation networks that often result from analyses of time series may themselves be best conceptualized as time-dependent. Overall the various sources of uncertainty documented here underscore the importance of transparent analysis, as well as the need for humility when drawing scientific conclusions.

## Methods

### Benchmarking statistical tests

We determined the true and false positive rates of various correlation tests by applying them to data from simulated ecosystems of two interacting species *x* and *y*. To calculate true positive rates, tests were presented with data from the same simulation. To calculate false positive rates, we presented each test with the *x* time series from one simulation and the *y* time series from an identical but independent simulation of the same ecosystem. True and false positive rates were calculated as the percentage of detections (*p ≤* 0.05) obtained by tests out of 1000 total pairs of *x* and *y* time series.

Based on their false positive rates, tests were considered either “ideal” (false positive rate *≤* 6.2%), “questionable” (6.2% to 10%) or “unacceptable” (*≥* 10%), as shown in, e.g., Fig 1C and 1F. We set 6.2% rather than 5% as the threshold for an ideal test to account for random fluctuations. More precisely, for a perfectly calibrated test with a true false positive rate of 5%, the number of false positives in 1000 trials follows a binomial distribution centered around 50. By setting our cutoff at 62 (i.e. 6.2% of 1000), we ensure that the probability of observing more than false positives than the cutoff is less than or equal to 5%.

For surrogate data tests, we first computed the correlation between the original *x* and *y*, and then compared this to a null distribution made from correlations between *x* and various *y*\* surrogates. In Fig 4, we also performed the reverse test, where the null distribution was made from correlations between *y* and *x*\* surrogates. For all surrogate methods except truncated time shift, the *p*-value was calculated as (1 + *n*_*larger*_)/(1 + *n*_*surr*_) where *n*_*larger*_ is the number of surrogate correlations that are as large as or larger than the original correlation, and *n*_*surr*_ is the total number of surrogate correlations. For surrogates produced by the random shuffle, stationary bootstrap, random phase, and twin methods, we used *n*_*surr*_ = 99 surrogates. For the circular time shift method, we used all possible surrogates (see Appendix). For the truncated time shift method, the number of surrogates and procedure to calculate the *p*-value are described in Appendix.

We set up surrogate data tests by pairing each of the 5 correlation statistics described in Table 2 with each of the 6 surrogate data procedures described in Table 1. See Appendix for a detailed introduction to each statistic and procedure.

For 3 correlation statistics (Pearson’s correlation coefficient, local similarity analysis, and Granger causality), we also assessed an analytical test–where the null distribution is obtained directly from a derived equation, rather than from surrogates. In the Pearson correlation test, the sample Pearson correlation coefficient is transformed into a *t*-statistic, whose null distribution is a *t*-distribution with an adjusted degrees of freedom based on the amount of autocorrelation in the time series [7]. For the analytical local similarity test, we estimate the “long-run variance” of the element-wise product of the two time series, and use that estimate to find a theoretical approximation for the *p*-value [26]. The analytical test for Granger causality was based on an *F*-test [27]. More detailed descriptions of these tests can be found in the Appendix.

All the above testing procedures are designed to test the null hypothesis that the two time series are independent, except for the analytical Granger causality test, which tests the null hypothesis of “Granger non-causality” (i.e. that the past of *x* does not contain unique information about the future of *y* [16, 17]). However, this exception is inconsequential for us since all of our independent cases are also Granger non-causal, and all of our dependent cases are constructed to be Granger-causal in both directions.

### Simulating two-species ecosystems

For each benchmark, we simulate the population dynamics in a pair of interacting species by integrating a system of ordinary differential equations over time steps of Δ*t* = 0.05 using the RK45 differential equations solver implemented in the package Scipy[41] and adding random process noise–representing population fluctuations such as from migration or stochastic environmental perturbations–to each variable between integration steps:

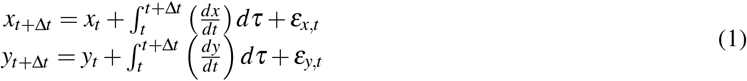

where *x*_*t*_ and *y*_*t*_ are the population density values of two species, *ε*_*x,t*_; *ε*_*y,t*_ are random variables, and Δ*t* = 0.05 is the time between instances of process noise. Since negative biomass is impossible, if a process noise term brought the population density of a species below zero, our simulation simply set the density to zero. For each simulation, functional forms for the rates of change (*dx*/*dt*; *dy*/*dt*) are depicted in the relevant figure, except for Fig 2, where the equations are

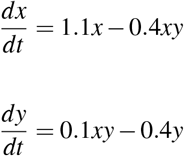

We start collecting time points after 3000 integration steps. This gives the system time to relax from its initial conditions and reach equilibrium, reducing dependence of the results on initial conditions. Then, we record observations of *x* and *y* every *n*_*s*_ integration steps, where *n*_*s*_ is typically 5 (which means the time between sampling points is *n*_*s*_Δ*t* = 0.25). In figures, time points are indexed with the variable *i*, where *i* = (*t* − 150)/(*n*_*s*_Δ*t*). For simulations with relatively smooth dynamics, we add measurement noise to each time point after simulation, which prevents numerical errors when computing correlation statistics. The measurement noise is drawn from a normal distribution centered at 0, with a variance chosen based on the stable population densities (e.g. 10^−2^ for the prey-predator system or 10^−6^ for competition). The details of these parameters on a per-simulation basis are given in Table 3.

## Notes

### Competing Interest Statement

The authors have declared no competing interest.

