## Appendix for "Subtle methodological variations substantially impact correlation test results in ecological time series"

### Statistics

#### Pearson correlation

Pearson correlation ($\rho$) is a widely used global measure of correlation. We calculate sample Pearson correlation $\rho$ for $N$ pairs of $x_i$ and $y_i$ ($1 \leq i \leq N$) with the following formula:

$$\rho = \frac{\sum_{i=1}^{N} (x_i - \bar{x})(y_i - \bar{y})}{\sqrt{\left[\sum_{i=1}^{N} (x_i - \bar{x})^2 \sum_{i=1}^{N} (y_i - \bar{y})^2\right]}}$$

where $\bar{x}$ and $\bar{y}$ are the sample means of $x$ and $y$ respectively. A positive value of $\rho$ indicates positive correlation, a negative value indicates negative correlation (anticorrelation), and a value of zero indicates no correlation. This statistic has no user-defined parameters.

In our tests based on surrogate data, we use the absolute value of the correlation coefficient $|\rho|$ as a statistic to test for independence between two time series. This allows for detection of either strong correlation or strong anticorrelation via a one-tailed test.

#### Local similarity analysis

Although Pearson correlation is well-suited for detecting global associations, it may have less success in identifying local associations (i.e. if $x$ is associated with $y$ in some but not all time points). Local similarity analysis (LSA) was designed to account for such situations [21] and has performed well in past simulation benchmarks using microbial time series data [28]. The algorithm to calculate this statistic for a pair of variables $x, y$ with measurement length $n$ is as follows:

1. Normalize $x$ and $y$ by ranking their values and applying an inverse cumulative distribution function of a standard normal distribution. This sets both variables to follow a standard normal distribution. For example, if the time series has 40 numbers, then the top number (belonging to the top 1/40=2.5%) will be assigned a value of 1.96. The normalized time series are denoted $\hat{x}$ and $\hat{y}$.

2. Initialize a positive association vector $S^+$ and a negative association vector $S^-$, with the initial values being zero. $S_0^+ = S_0^- = 0$.

3. For $i \in \{1, 2, ..., n\}$ where $n$ is the length of the series:

$$S_i^+ = max[0, S_{i-1}^+ + \hat{x}_i * \hat{y}_i]$$

$$S_i^- = max[0, S_{i-1}^- - \hat{x}_i * \hat{y}_i]$$

   where $\hat{x}_i * \hat{y}_i$ is the $i$th element of $\hat{x}$ multiplied by the $i$th element of $\hat{y}$. Note that if $x_i$ is greater than the mean of $x$, then $\hat{x}_i$ will be positive. Thus, if both $x_i$ and $y_i$ are greater than their respective means or if both are less than their respective means (as in a positive association), then $\hat{x}_i * \hat{y}_i$ will be positive. This will increase the value of $S_i^+$, and decrease the value of $S_i^-$ (as long as $S_{i-1}^-$ is nonzero). If $\hat{x}_i * \hat{y}_i$ is negative, the value of $S_i^-$ increases and the value of $S_i^+$ decreases (as long as $S_{i-1}^+$ is nonzero).

4. Calculate local similarity $LS = max(S_i^+, S_i^-)/n$. Take the maximum value from $S_i^+$ and $S_i^-$, and divide it by the measurement length $n$ to obtain the statistic. Note that $LS$ is nonnegative here, enabling detection either strong correlation or strong anticorrelation via a one-tailed test, similar to the set up with Pearson correlation. In a two-tailed test, local similarity from negative associations would be represented by a negative number.

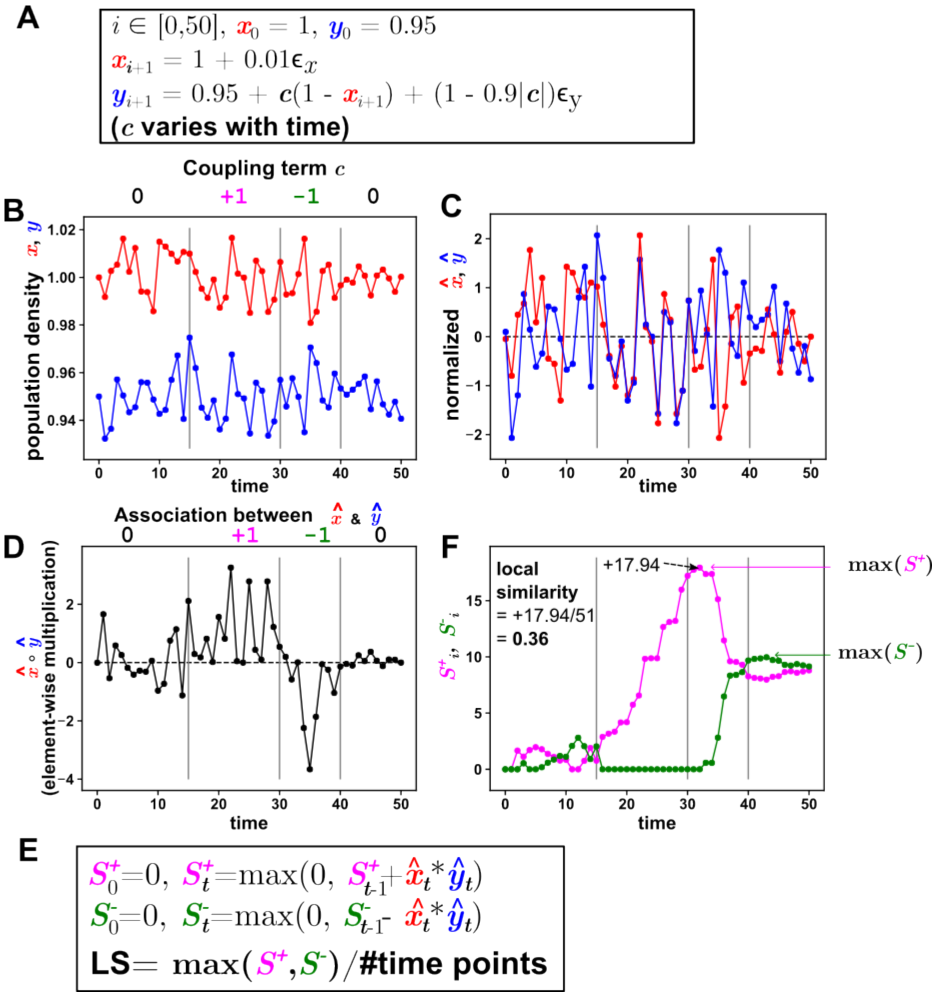


Figure 1: Local similarity analysis looks for transient correlations between time series. (A) A system of two variables with coupling coefficient $c$ that varies over time. Coupling can be positive, negative, or zero. (B) The two variables measured over 51 time points. Time intervals are labeled with their corresponding coupling values. (C) Both time series are scaled to a standard normal distribution. (D) The elementwise product of the normalized time series. This vector tends to be positive when the time series are positively associated, and negative when the time series are negatively associated. (E) Definitions of $S_i^+$ and $S_i^-$. For example, if $\hat{x}_i * \hat{y}_i$ is positive, then $S_i^+$ increases from $S_{i-1}^+$ and $S_i^-$ decreases from $S_{i-1}^-$ (with the minimal value of each being zero). The Local Similarity score is calculated as the following: take the maximum value of all elements from both vectors and divide it by the length of the time series. (F) Plots of $S^+$ and $S^-$ over time, and the resulting local similarity.

#### Granger causality

A variable $y$ is said to Granger-cause another variable $x$ if a model incorporating the history of all variables predicts the future of $x$ more accurately than a similar model that excludes the history of $y$ [12, 13, 4]. We use linear Granger causality, where we predict the future of $x$ using linear regression on the history of $x$ and $y$ (Fig 2). We quantify Granger causality as the normalized difference between the errors of the two models, which should be strictly non-negative and approximately 0 for time series with no causal relationship. Specifically, to calculate the Granger causality of $y$ on $x$ in a two-variable system with sample times $i = [0, 1, ..., N-1]$, we build two linear autoregressive models: a "full" model including information about the history of $x$ and $y$, and a "base" model with only the history of $x$.

$$x_i = \sum_{j=1}^{p} [\beta_j^{full} x_{i-j} + \alpha_j^{full} y_{i-j}] + \gamma^{full} + \varepsilon_i^{full}, i \in [p, N-1]$$

$$x_i = \sum_{j=1}^{p} \beta_j^{base} x_{i-j} + \gamma^{base} + \varepsilon_i^{base}, i \in [p, N-1]$$

Here, the hyperparameter $p$ is the length of the history vectors. We scanned $p = 1, 2, ..., 15$, and selected the value of $p$ that gave the model the highest Akaike information criterion (AIC), a measure of model quality. The key Granger causality statistic is the normalized difference between two models' errors, quantified as an $F$-statistic. We use the implementation in the Python package statsmodels [24].

$$F = \left(\frac{N-3p-1}{p}\right) \frac{\sum_{i=p}^{N-1} [\varepsilon_i^{base}]^2 - \sum_{i=p}^{N-1} [\varepsilon_i^{full}]^2}{\sum_{i=p}^{N-1} [\varepsilon_i^{full}]^2}$$

The second half of this expression represents the difference between the sum of squared residual of the base model and that of the full model normalized against the full model. The first half of this expression is given by $d^{full}/(d^{full} - d^{base})$ where $d^{base}$ and $d^{full}$ are the numbers of degrees of freedom in the base and full models respectively. Here, the number of degrees of freedom for each model is simply given by the number of time points minus the number of parameters. Specifically, for the base model, there are $N - p$ time points with $p + 1$ parameters, and hence $d^{base} = N - 2p - 1$. Similarly, since the full model has $2p + 1$ parameters, $d^{full} = N - 3p - 1$.

In a parametric test, if $y$ does not Granger-cause $x$, then we would expect the Granger causality statistic of $y$ on $x$ to follow an $F(d^{full} - d^{base}, d^{full})$ distribution, and so we may reject the null hypothesis of Granger non-causality on the basis of a one-tailed $F$-test. This is not the same as saying that $x$ and $y$ are independent. Rather, this procedure tests "conditional independence", independence between later values of $x$ and $y$ conditioned on earlier values of $x$. Technically, it isn't appropriate to use the Granger causality framework with our collection of surrogate data methods, which are designed for the null hypothesis of independence rather than conditional independence [25]. However, since Granger causality has been previously used with similar surrogate data methods [8, 18, 17], we benchmark its performance here for comparison.

#### Cross map skill

Cross map skill is a statistic that measures how well one time series can be predicted using another [26]. It is commonly used in procedures that attempt to determine the direction of causation [26, 30], but we assess it within the more modest context of detecting dependence.

Cross map skill works as follows (Fig 3). To predict $y_k$ from $x$ (Fig 3A, Step 1), one decides on parameters $D$ and $\tau$ for a "delay vector", which is a segment of $x$'s history with dimension $D$ (in the figure $D = 3$) and a delay spacing $\tau$ (in the figure $\tau = 1$). The delay vector is constructed from data points $x_k, x_{k-\tau}, x_{k-2\tau}, ..., x_{k-(D-1)\tau}$ (Fig 3A, Step 2). One then looks up $D + 1$ delay vectors in the history of $x$ that most resemble this segment, called "neighbors" (Fig 3A, Step 3; Fig 3C). The next step is to look up each corresponding contemporaneous $y$ value (Fig 3A; Step 4), and weigh them according to how similar the neighbor $x$ delay vector is to the original $x$ delay vector (measured as Euclidean distance). One can then predict the $y$ as the weighted average of these "neighbor $y$ values" (Fig 3A; Step 5). The cross map skill is the Pearson correlation $\rho$ between each true value of $y_k$ and the predicted value of $y_k$ (Fig 3B).

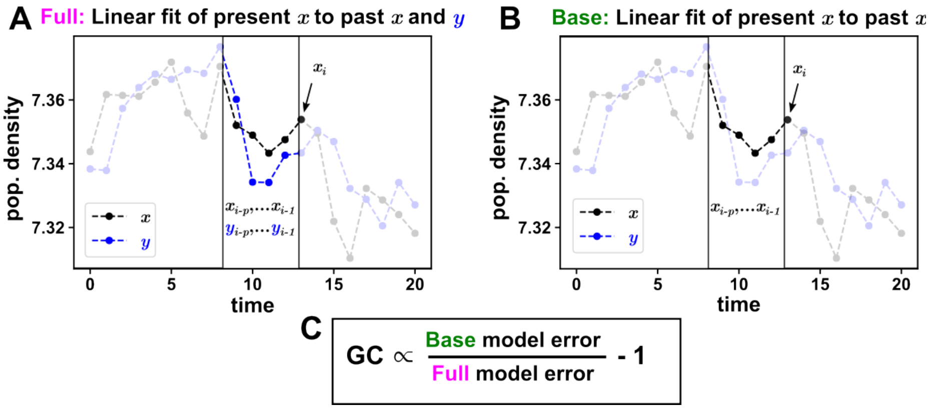


Figure 2: Testing whether $y$ Granger causes $x$. (A) Full model: We construct a linear model of the target variable $x_i$ as a function of a vector of past $x$ values $(x_{i-1}, x_{i-2}, \ldots x_{i-p})$ and a vector of past $y$ values $(y_{i-1}, y_{i-2}, \ldots y_{i-p})$. We use ordinary least squares to estimate the coefficients of this model and calculate its error as the sum of the squared residuals. (B) Base model: we remove the past values of $y$ from the model. In the same way, we fit the model and calculate its error. (C) The Granger causality statistic is the normalized difference between the (sum of squared) error of the full model using $y$ and the base model not using $y$.

#### Mutual information

Mutual information is a nonlinear measure of dependence between variables. We use Kraskov's $k$-nearest neighbors estimator [15] of mutual information, summarized as follows.

Let the length-$N$ time series be a set of two-element vectors $S_i = (x_i, y_i)$, $i \in [1, N]$. For each time point $i$, we first find $S_i$'s $k$-nearest neighbor vector $S_u$. For example, if $k = 1$, we find the vector closest to our vector of interest, and if $k = 3$, we find the third closest vector. To quantify the distance between $S_i = (x_i, y_i)$ and $S_u = (x_u, y_u)$, we use $max(|x_i - x_u|, |y_i - y_u|) = \varepsilon_i/2$. Then, we find $n_x(i)$, which is the number of values of $x$ such that $x_i - \varepsilon_i/2 < x < x_i + \varepsilon_i/2$, not including $x_i$. We find $n_y(i)$ in a similar way: the number of values of $y$ such that $y_i - \varepsilon_i/2 < y < y_i + \varepsilon_i/2$, not including $y_i$. Then, for $X = (x_1, \ldots, x_N), Y = (y_1, \ldots, y_N)$, we can estimate the mutual information $I(X, Y)$ as follows:

$$\hat{I}(X,Y) = \psi(N) + \psi(k) - \frac{1}{N}\sum_{i=1}^{N}[\psi(n_x(i)+1) + \psi(n_y(i)+1)]$$

where $\psi$ is the digamma function.

### Surrogate methods

#### Random shuffle

In the random shuffle method, also known as the permutation method, one generates a surrogate time series of by shuffling the original time series (Fig 5A). This maintains the original distribution but destroys any temporal structure in the original data. Its associated modeling assumption is that the original data can be described by independent and identically distributed random variables [16]. This null hypothesis is commonly violated by time series data, since a system's future often depends on its past. Despite its propensity for a high false positive rate when applied to time series, the permutation test is commonly used.

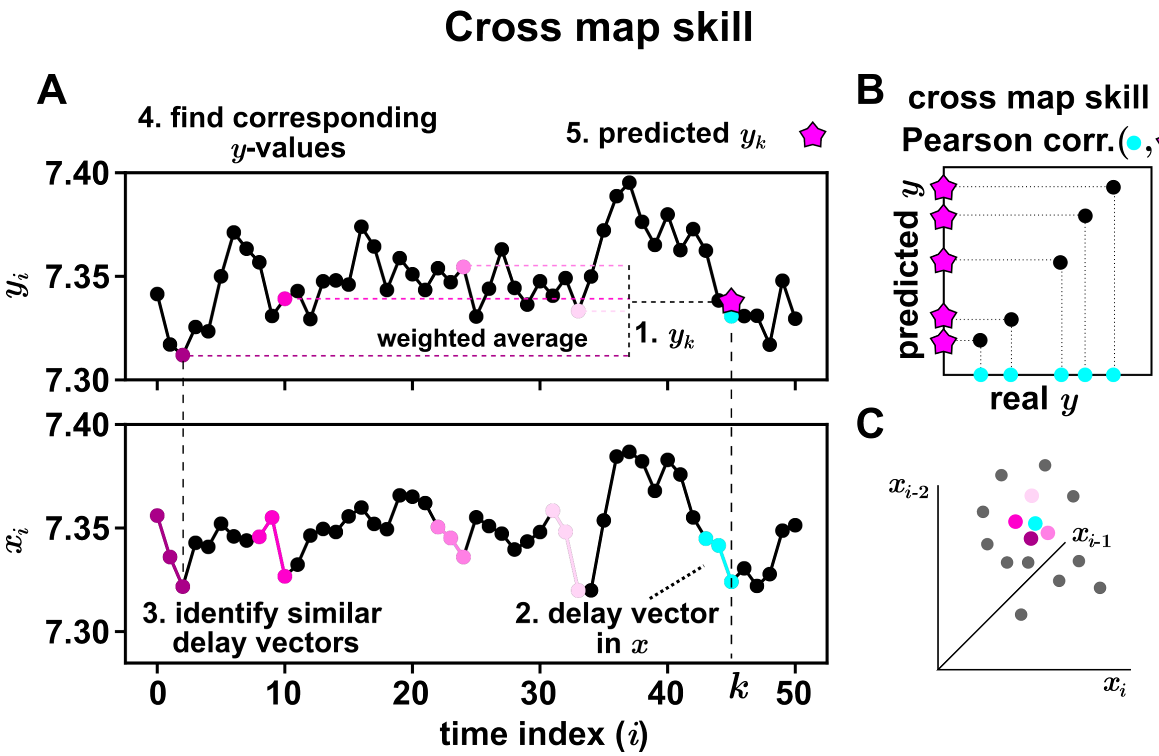


Figure 3: Cross map skill is a measure that quantifies the amount of information about one variable stored in the history of another. (A) We want to predict observations of $y_k$ (Step 1) with corresponding delay vectors from $x$ (Step 2). A delay vector is made of equally spaced time points from a time series. Here, the $x$ delay vector corresponding to $y_k$ - the focal vector - is colored in cyan. We find some delay vectors in $x$ (magenta with various shades) that are nearby in Euclidean space to the focal vector (Step 3; Panel C) and find their corresponding $y$ values (Step 4). Then, we average these $y$ values, weighted by their similarity to the focal vector, to predict $y_k$ (Step 5). (B) We repeat (A) many times for different time points $k$. The cross map skill is the Pearson correlation between the predicted $y$-values and their true values. (C) A scatter plot of delay vectors in the delay space.

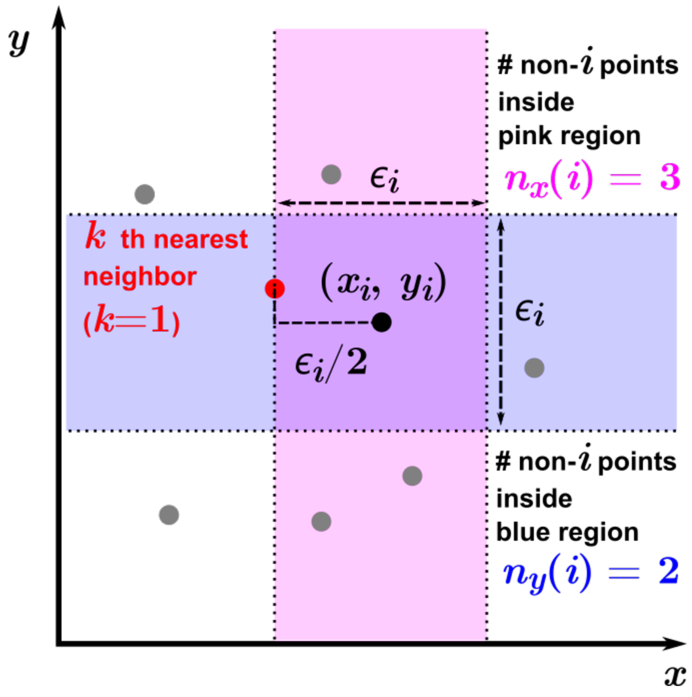


Figure 4: Mutual information is a statistic that quantifies dependence. We use an estimator based on nearest neighbors to estimate mutual information between two time series. For each time point $i$ in the time series, we find the point with the $k$th shortest "maximum distance" from the point $(x_i, y_i)$, where maximum distance is the greatest difference along any coordinate dimension. We call this distance $\varepsilon_i/2$. Then, we count the number of data points in $x$ that are less than $\varepsilon_i/2$ away from $x_i$ (inside the pink region), and the number of data points in $y$ that are less than $\varepsilon_i/2$ away from $y_i$ (inside the blue region). These counts are used to estimate the mutual information between $x$ and $y$.

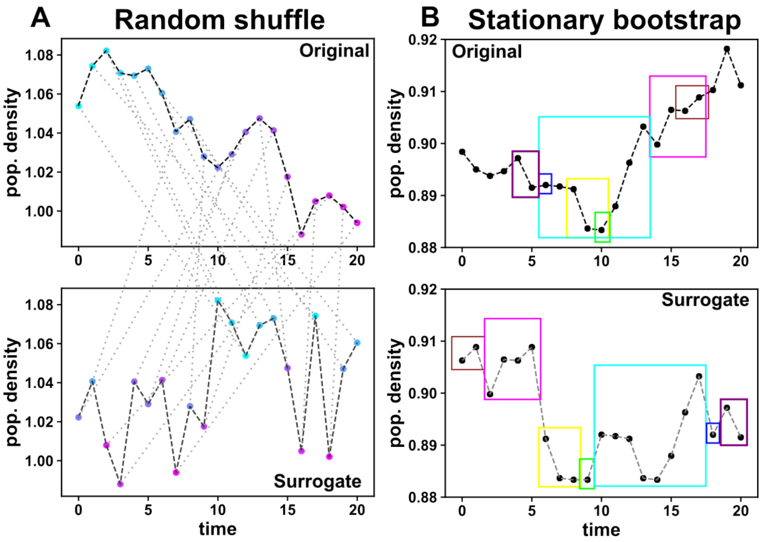


Figure 5: Permutation-based surrogate data methods. (A) The permutation or the random shuffle method shuffles the positions of data points in the time series. (B) The stationary bootstrap surrogate data method samples blocks of consecutive data points from the original time series with resampling. The expected size of each block is $1/\alpha$, where $\alpha$ is a user-defined parameter (hyperparameter). In this study, we select $\alpha = 0.05$.

#### Stationary bootstrap

The stationary bootstrap method [19] partially preserves autocorrelation while still somewhat scrambling the original data. Here, instead of simply shuffling individual time points, one can randomly select consecutive blocks of data to construct a surrogate. The method is as follows:

Consider a time series $x_i, i \in [1,N]$. The goal is to construct a surrogate time series $x_i^* = x_{j(i)}, j \in [1,N]$ with the same length.

1. Choose a random integer $j(1) \in [1,N]$. We set $x_1^* = x_{j(1)}$.
2. For $i \in [2,N]$, set $x_i^* = x_{j(i)}$ where
   (a) With probability $1-\alpha$, $j = j(i) = (j(i-1)+1) \mod N$. That is, go to the next point in the original time series.
   (b) With probability $\alpha$, pick a new random integer $j(i) \in [1,N]$. That is, jump to another place in the original time series.

Thus, the surrogate time series is made of randomly sized and shuffled blocks of data with average length $1/\alpha$ (Fig 5). For this study, we set $\alpha = 0.05$.

Although shuffling in blocks may not be as disruptive as the complete random shuffle described just previously, this block shuffling method still somewhat disrupts temporal properties of the original sequence by creating discontinuity between blocks. Indeed, a Granger causality test based on this method is said to be inexact [8].

#### Fourier transform-based surrogate methods

Fourier transform-based (also known as random phase) surrogate data methods preserve the power spectrum of the data (i.e. amplitude against frequency), which also approximately preserves its autocorrelation [6]. In this class of methods, one produces surrogate data by decomposing the original data into a set of sinusoidal waves with a Fourier transform, shift each wave component by a randomized phase $\phi \in [0, 2\pi)$, and add the shifted waves together to get the surrogate data (Figure 6). This method, called the random phase or Fourier transform (FT) method, works for the null hypothesis that the original data comes from a stationary linear Gaussian process that is independent of the other time series.

The random phase method can be adjusted to work for nonlinearly *rescaled* linear Gaussian processes. An example of this is a measurement $x_t = (a_t)^3$ where $a_t$ is a stationary linear Gaussian process. While $x_t$ itself is not linear, it can be made linear with an invertible, time-independent scaling function. Although in this example, we know how to invert the nonlinear $x_t$ to $a_t$ (by taking 1/3 power), we generally do not know the inversion function. In this case, we can use the amplitude-adjusted Fourier transform (AAFT) method. Specifically, one performs the same phase-randomizing procedure, but preprocesses the data by rescaling it to follow a Gaussian distribution (assigning each original number according to its rank with a corresponding number from a random Gaussian time series). Then, we perform random phase procedure on the rescaled data, and obtain a series of numbers over time. For each number, based on its rank, we assign it the value corresponding to the original value of the same rank to get the final surrogate data. Because of this preprocessing, surrogate data from the AAFT method might not have the exact same power spectrum as the original data. The iterative amplitude-adjusted Fourier transform (IAAFT) method accounts for this mismatch by iterating between a step that adjusts the Fourier amplitudes and a step that applies a nonlinear rescaling, so that after a sufficient number of steps, surrogates have both an accurate power spectrum and an accurate distribution [22, 23, 16].

Fourier transform-based surrogate methods are very commonly used for hypothesis testing in time series in fields such as geophysics and neuroscience, although they are less commonly used in microbiome studies [14, 11, 5].

In this study, we use the IAAFT method as defined in the Python package pyunicorn [9]. We set the maximum number of iterations to 200.

#### Circular time shift

Time-shifting surrogate data methods are some of the least computationally expensive ways to generate a null distribution, and preserve most properties of the original data. We generate circular time shift surrogates by temporally shifting each element of a time series, and wrapping any points that "fall off the end" back to the beginning [3] (Fig. 8A). For example, if the original time series is $\{1,2,3,4\}$, then this method would produce three surrogates: $\{2,3,4,1\}$, $\{3,4,1,2\}$, and $\{4,1,2,3\}$.

#### Pre-processing periodic time series for use with Fourier transform or time shift methods

The circular time shift involves a wrap-around step for obvious reasons, and the stationary bootstrap method can also involve a wrap-around step if the surrogate reaches past the end of the original time series. When the beginning and end of a time series are very different, this can introduce artifacts (Fig 7A). Fourier-based surrogates can also suffer because the Fourier transform assumes that a time series is periodic and has a length that is an integer multiple of the period. To help address these problems, one can trim the time series to make the ends match (Fig. 7 B). We use the trimming method described in [16]. This method takes a parameter $p$ as the length of time segments to match at the beginning and end of the trimmed time series. We estimate the period of the time series $T$ with a fast Fourier transform, and set the parameter $p = T/10$. Lancaster et al. [16] advocated this procedure particularly for periodic processes, but we suspect that the benefits extend to other systems, and thus we use the trimming procedure for all systems in our benchmark case studies when using circular timeshift and random phase tests. Since the stationary bootstrap method already suffers from many discontinuities, we did not use the trimming procedure with that method.

#### Truncated time shift

One way to apply the time-shift method without any need for preprocessing and avoid discontinuities is to truncate the time series and take time shifted subsets of original data (Fig8B). Here, the "original" data set from which we calculate the observed statistic would be a subset of the actual time series. and the surrogate data would be a different subset from the same time series with the same length [3]. This method is valid for testing the null hypothesis that two time series $x$ and $y$ are independent, if at least one of the time series (e.g. $y$) is stationary

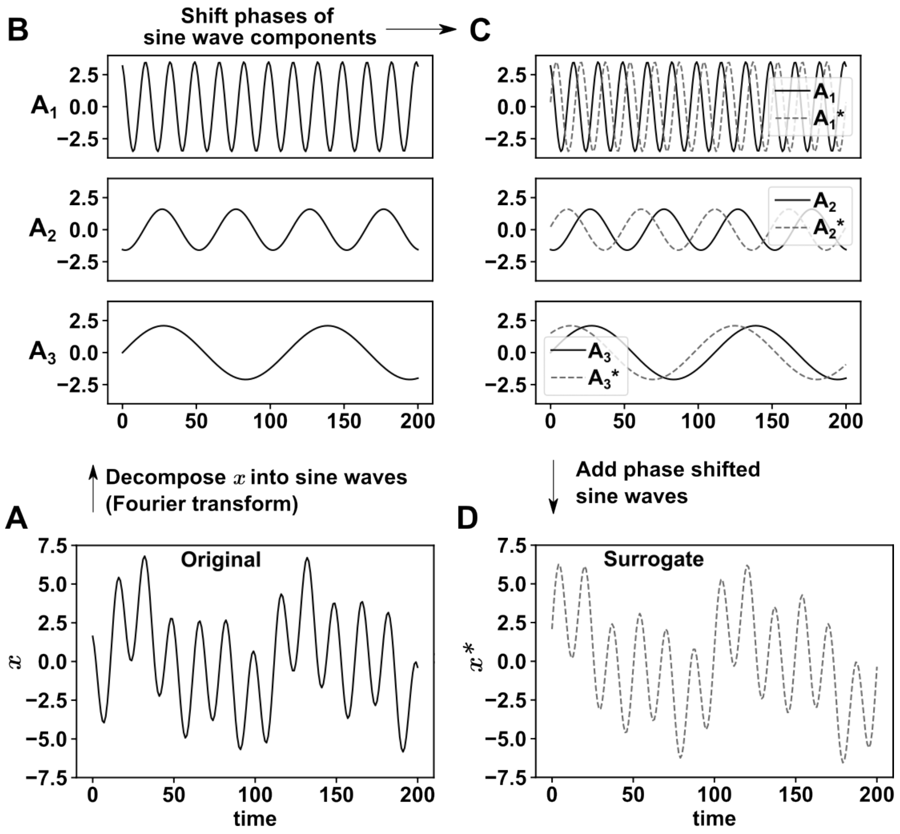


Figure 6: The basic “random phase” or “Fourier transform” surrogate method involves decomposing the time series into a set of sine waves (A to B), changing the phase of each wave by a random value (C), and adding the resulting waves back together to get the final surrogate time series (D).

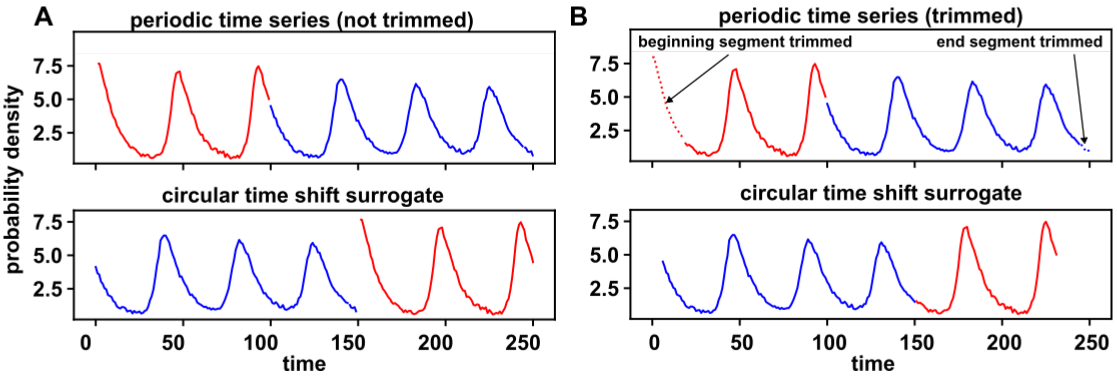


Figure 7: Creating a circular time shift surrogate from a periodic time series without or with trimming. (A) Without trimming, a discontinuity artifact is created. (B) The discontinuity artifact is alleviated by trimming the time series to match the two ends. ) A grid search algorithm chooses a segment from each end of the time series to remove (dotted lines), such that the ends of the remaining time series have similar values.

[16, 18, 31]. The stationary series $y$ is used to generate surrogate $y^*$ and correlation between $x^{trunc}$ and $y^*$ is then compared to those between $x$ and $y^*$.

Here, we use truncated time series of length $N-2r$ and use subsets of this length as surrogate time series. We select the largest possible $r$ based on the length of the time series $N$ such that $r+1 < \frac{N}{4}$ and $\frac{r+1}{20}$ is a positive integer. For a time series indexed from 0 to $N$, the "true" time series is considered to be the subset at the index range $[r, N-r-1]$. The "surrogate" time series are sampled from the range $[r+\delta, N-r-1+\delta]$ where $\delta \in [-r, +r]$.

The original truncated time shift method has been shown to display false positive rates above the significance cutoff, because of the dependence between surrogates (e.g. two neighboring surrogates share the vast majority of data). To guarantee a false positive rate at or below the significance level, we use an adjusted $p$-value calculated as $(N_{\geq}+1)/(r+1)$ (where $N_{\geq}$ is the number of shifted correlations that are at least as large as the unshifted correlation) [31].

**Twin method**

The twin surrogate method was designed to test for phase synchronization between two oscillating time series [16, 27]. In this method, we embed (replot) data in a delay space (Figure 9B). "Neighbors" of a focal point are all the points within the predefined distance to the focal point. Two points in the delay space are "twins" if they have the same set of neighbors (Figure 9C). To make surrogate data, a random starting point is chosen from the original time series, and the next point will depend on whether the current point has twins. If no twins, then the next point will be the following point from the current point (or a randomly chosen point if the current point is at the end of the time series). If the current point has twins, then the next point can be the following point from one of the twins (Figure 9D). As a result, the surrogate series jumps around in time between similar system states. This method seems to roughly correspond to a null hypothesis that two time series are independent and Markovian (i.e. what happens next only depends on the current time point's delay vector).

When applying the twin surrogate method, we choose embedding dimension and the delay from a grid search (both in range [1,8]) and choose the values that give the highest cross map skill of $y$ to itself (see the section on convergent cross mapping). Neighbor distance threshold is chosen automatically so that on average, each point in the delay space has a specified number of neighbors (here, 10% of total).

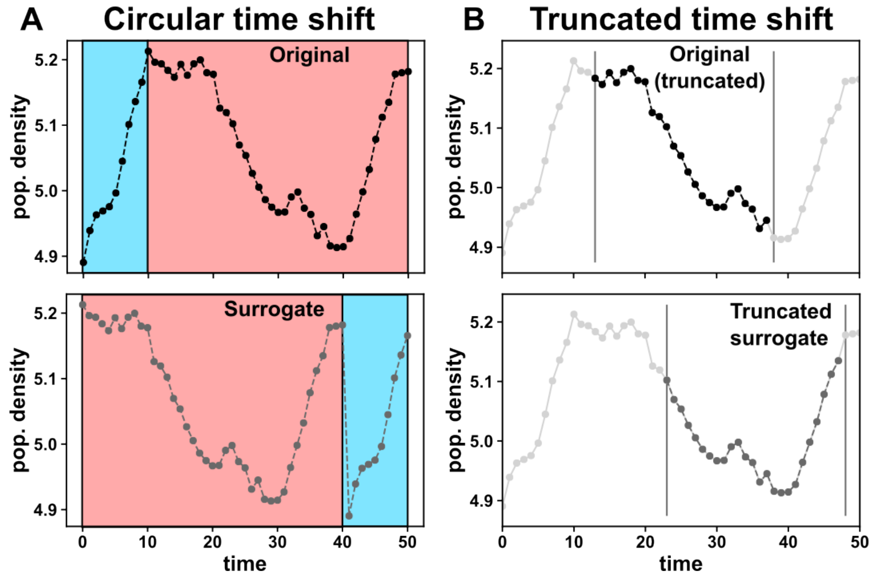


Figure 8: Time shift-based surrogate methods. (A) Circular time shift consists of shifting a time series in time by a random amount (red) and the displaced original data (blue) are stitched back at the end. This can create a discontinuity (dotted line). (B) To avoid the problem of discontinuity, one can truncate the original time series and use other parts of a time series as surrogates.

### Analytical tests

#### Parametric autocorrelation-aware hypothesis test with Pearson correlation

We wish to benchmark the surrogate data methods against an analytical method of detecting dependence between time series. Multiple methods exist for estimating the sample variance of Pearson correlation between independent autocorrelated time series [2, 7, 1, 10]. See [20] for further discussions. Here, we use an adjusted $t$-test from Clifford et al. [7] that accounts for autocorrelation in the time series due to its superior performance compared to three other alternative tests [31]. Since this method is completely analytical, it is much less computationally expensive than the surrogate data methods described in this paper.

To test if two time series $x, y$ with length $N$ are independent, we first calculate their Pearson correlation $\rho_{xy}$ (see "Statistics" section). The next step is to analytically derive the null distribution of $\rho_{xy}$ under the assumption that the time series are independent.

We define autocovariance matrices $\Sigma_x, \Sigma_y$ such that $\Sigma_x$ at index $ij$ is the covariance between $x$ at time $i$ and $x$ at time $j$. If both time series are covariance stationary, then $cov(x_1, x_2)$ will be the same as $cov(x_2, x_3), cov(x_3, x_4)$, and so on. Thus, the estimated autocovariance matrices will have equal values along the diagonals. It will also be symmetric. Some researchers will weight autocovariance terms such that shorter lags have greater "weight", with lags past a certain threshold being ignored entirely [20]. Here, all lags greater than 25% of the time series length are ignored.

With these matrices approximated, we find the sample variance of the Pearson correlation between two independent time series (see [10] for derivations) :

$$var(\hat{\rho}_{xy}) = \frac{tr(B\Sigma_x B\Sigma_y)}{tr(B\Sigma_x)tr(B\Sigma_y)}$$

Here, $B = (I_N - N^{-1}J_N)/N$ where $I_N$ is the $N$ by $N$ identity matrix, and $J_N$ is the $N$ by $N$ matrix of all ones, and $tr()$ - the trace of a matrix - is the sum of its diagonal terms. If this variance term is negative, we set it to the reciprocal of the time series length.

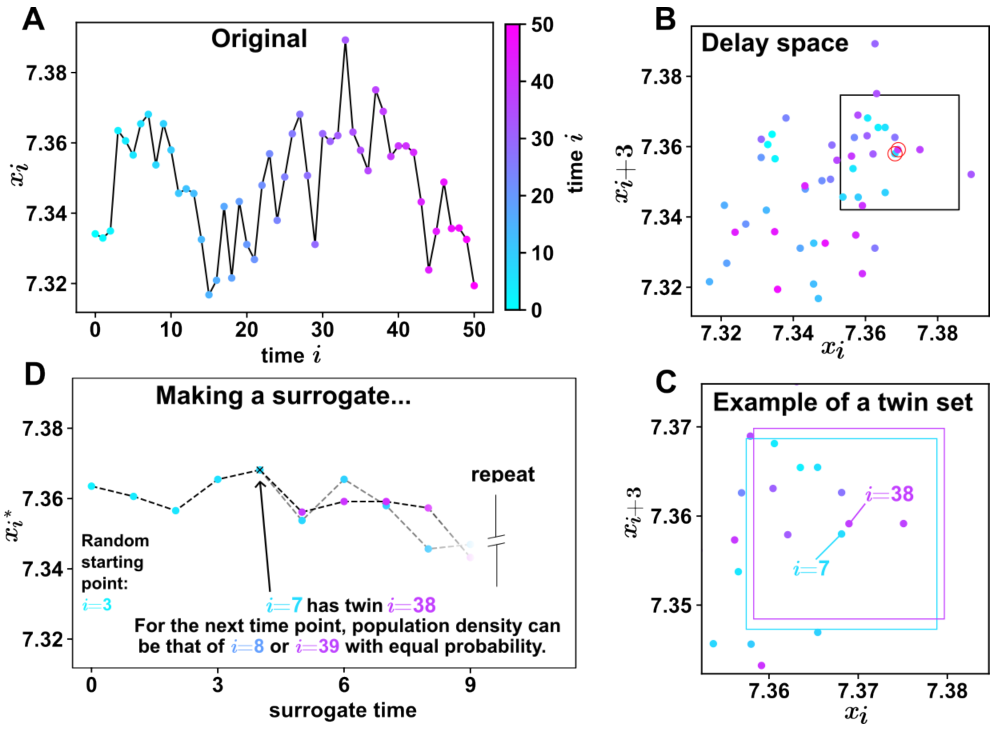


Figure 9: The twin method of constructing surrogate time series. (A) The original time series, with each point colored by time. (B) A delay space of delay vectors. A delay vector consists of equally spaced time points, and the user can select the length of the vector ("embedding dimension") and the time delay spacing between elements. Here, we use embedding dimension of 2 and a delay of 3 (i.e. a delay vector is $(x_i, x_{i+3})$. (C) A delay vector's neighborhood is the set of delay vectors within a certain maximum distance of the focal vector. The neighborhoods of delay vectors at $i = 7$ and $i = 38$ are shown as colored squares around the points. Delay vectors are twins if they have the same neighborhood, as is the case for delay vectors at $i = 7$ and $i = 38$. (D) To construct the surrogate time series, we start by randomly selecting a point from the original time series. If the delay vector at this point has no twins, the next point will be the following time point in the original series. Otherwise, the next point will be the following point from a randomly chosen twin.

Finally, we define the effective sample size $n_{eff} = 1 + [var(\hat{\rho}_{xy})]^{-1}$. We use the effective sample size to find the Student's t-statistic.

$$t_{xy} = \frac{\rho_{xy}\sqrt{n_{eff} - 2}}{\sqrt{1 - r_{xy}^2}}$$

The null distribution is a $t$-distribution with mean of 0, variance of 1, and $n_{eff} - 2$ degrees of freedom. We perform a two-tailed test: if $t_{xy}$ is in the top 2.5% or bottom 2.5% of the null distribution, we reject the null hypothesis.

##### Significance test for local similarity

The original paper on LSA [21] tested for statistical significance using a permutation test, which is generally inappropriate for time series. Since then, a variety of improvements have been made. Here, we use the test of Zhang et al. [32]. This method can handle autocorrelation, so long as both time series are stationary.

To apply the method, let $x_i$ and $y_i$ be two time series ( $i$=1,..., $N$). We normalize both time series to have a mean of zero: $X_i = x_i - \bar{x}, Y_i = y_i - \bar{y}$. We define the autocovariance function of $X_i$ at lag $k$ as follows:

$$\hat{\gamma}_X(k) = \frac{1}{N} \sum_{j=1}^{N-|k|} (X_{j+|k|} - \bar{X})(X_j - \bar{X}), k \in [1, N-1]$$

We can define $\hat{\gamma}_Y(k)$ analogously. With these functions defined, we can estimate the long-run variance of the elementwise product of $X$ and $Y$ as follows:

$$\hat{\omega}^2 = \hat{\gamma}_X(0)\hat{\gamma}_Y(0) + 2\sum_{k=1}^{b_w}(1 - \frac{k}{b_w})\hat{\gamma}_X(k)\hat{\gamma}_Y(k)$$

$b_w$ is a bandwidth term used to weigh autocorrelation between "nearby" lags with higher value than autocorrelation between "distant" lags. Its formula can be found in [32].

Finally, we plug our estimate of $\hat{\omega}$ into the following equation, which theoretically approximates the probability of getting a certain local similarity score between two independent time series with similar size and autocorrelation:

$$p = P(\frac{LS}{\hat{\omega}\sqrt{N}} \geq a) = 1 - 8\left[\sum_{k=1}^{\infty}\left\{\frac{1}{a^2} + \frac{1}{(2k-1)^2\pi^2}\right\}\exp\left\{\frac{-(2k-1)^2\pi^2}{2a^2}\right\}\right]$$

where $N$ is the number of time points, $\sigma$ is the standard deviation of the elementwise product of the two normalized time series, and $a$ is the local similarity score between $x$ and $y$. This is a simplified version of the equation found in [29] because it does not account for delays.

##### Granger causality $F$-test

In some cases, we try to detect linear Granger causality with an $F$-test. We use the implementation of this test given by the function "grangercausalitytests" in the Python package statsmodels [24], which fits two ordinary least squares models with different sets of independent variables and gives the difference in residual errors between the two models as an $F$-statistic. The formula for the $F$-statistic is found in the "Statistics" section of this appendix.
